# PaintSHOP enables the interactive design of transcriptome- and genome-scale oligonucleotide FISH experiments

**DOI:** 10.1101/2020.07.05.188797

**Authors:** Elliot A. Hershberg, Jennie L. Close, Conor K. Camplisson, Sahar Attar, Ryan Chern, Yuzhen Liu, Shreeram Akilesh, Philip R. Nicovich, Brian J. Beliveau

## Abstract

Fluorescent *in situ* hybridization (FISH) allows researchers to visualize the spatial position and quantity of nucleic acids in fixed samples. Recently, considerable progress has been made in developing oligonucleotide (oligo)-based FISH methods. These methods have enabled researchers to study the three-dimensional organization of the genome at super-resolution and visualize the spatial patterns of gene expression for thousands of genes in individual cells. While considerable progress has been made in developing new molecular methods that harness complex oligo libraries for FISH, there are few existing computational tools to support the bioinformatics workflows necessary to carry out these experiments. Here, we introduce Paint Server and Homology Optimization Pipeline (PaintSHOP), an interactive platform for the reproducible design of oligo FISH experiments. PaintSHOP enables researchers to identify probes for their experimental targets efficiently, to incorporate additional necessary sequences such as primer pairs, and to easily generate standardized files documenting the design of their libraries. Our platform integrates a machine learning model that quantitatively predicts probe specificity on the genome scale into a dynamic web application that creates ready-to-order probe sets for a wide variety of applications. The goal of this freely available web resource is to democratize and standardize the process of designing complex probe sets for the oligo FISH community. PaintSHOP can be accessed at: paintshop.io

## Introduction

Fluorescent *in situ* hybridization (FISH) is a powerful technique that allows researchers to visualize the distribution of RNA and DNA at single-cell resolution in fixed samples. Since the introduction of *in situ* hybridization in 1969^1^ and the subsequent development of FISH^2–4^, the method has continued to be used, updated, and refined as new technologies became available. In particular, many improvements have been made to the process of constructing and labeling the probes used in a FISH assay. One such advance was the introduction of the nick translation method^5,6^, which increased the specific activity of labeling. Subsequent advances in DNA sequencing and synthesis technologies have spurred the development of a new generation of advanced FISH techniques that utilize oligonucleotide (oligo) libraries as a source of probe material.

Oligo FISH probes offer many advantages compared to conventional probes derived directly from genomic material. Synthetic oligos allow for greater engineering precision, as sequences can be optimized to have specific thermodynamic properties, bind to precisely defined targets, avoid repetitive sequences, and incorporate a variety of labeling schemes. Researchers have visualized multicopy targets such as repetitive DNA^7–9^ and mRNA^10–12^ using one to a few dozen individually synthesized oligos. Approaches have also been developed to leverage complex oligo libraries created by massively parallel array synthesis^13^ to perform oligo FISH experiments targeting single-copy chromosomal regions^14,15^. We have previously described Oligopaints^16^, a PCR-based method for the generation of highly specific and efficient RNA and DNA FISH probes from complex oligo libraries. The use of complex oligo libraries has enabled massively multiplexed DNA FISH experiments^17–20^ and spatial transcriptomics approaches targeting hundreds to thousands of individual mRNA molecules^21–23^.

While many experimental advances have been made using oligo FISH probes, comparatively little progress has been made in developing computational tools that support the design of these probes. Several computational tools exist for various related problems such as designing oligo probes against targets such as bacterial rRNA^24,25^, large pools of oligo pairs^26–29^, padlock probes^30,31^, or microarrays^32^.

Previously, we introduced OligoMiner^33^, a bioinformatic pipeline developed to address the bottleneck of computationally designing probe sets for Oligopaints and other oligo FISH approaches. Additionally, a bioinformatic resource called iFISH^34^ was created to provide access to pre-designed “probe spots” or to retrieve probes covering a single region. However, when it comes to supporting a wide degree of experimental designs and the necessary steps required to generate complete probe sets, these existing tools either require considerable bioinformatics expertise or lack the scalability and flexibility needed to complete the desired design workflows. To our knowledge, no framework exists to solve common problems such as appending additional necessary sequences to probe sequences like primer pairs for PCR amplification or building probe sets against multiple targets.

Here, we introduce Paint Server and Homology Optimization Pipeline (PaintSHOP), a platform that enables the interactive design of oligo-based FISH experiments at transcriptome- and genome-scale. PaintSHOP consists of two components: 1) a bioinformatic pipeline for large-scale thermodynamics modeling of probe specificity using machine learning, and 2) an interactive web application for the creation of ready-to-order oligo library designs against any target in the genome or transcriptome (paintshop.io). The predictions from the machine learning model are integrated directly into the web application, making the technology entirely accessible through the browser. The result is a freely available community resource that bridges the gap between probe set design and experiment.

## Results

### Homology Optimization Pipeline

While it is essential to identify as many candidates as possible when generating an oligo probe set, it is equally important to screen the candidates identified for the possibility of binding to regions in the genome other than their intended target. For

FISH, minimizing off-target binding is essential for successful visualization and interpretation of the spatial position of fluorescent signal measured. In order to model probe specificity, we developed a resource termed the ‘Homology Optimization Pipeline’ (**Fig. 1**). This pipeline integrates next-generation sequencing (NGS) alignment with machine learning into an automated and scalable Snakemake^35^ workflow to quantitatively predict the specificity of each probe candidate. In order to use the pipeline, users first need to input candidate probe sequences in FASTQ-format^36^. First, the pipeline uses the Bowtie2 NGS read aligner^37^ to align all probe candidates against the genome that they are targeting (**Fig. 2A**). Bowtie2 has a “speed/sensitivity/accuracy trade-off space” where there is a balance between how quickly the alignment can be performed, how accurate the alignments are, and how sensitive the algorithm is to all possible alignments. As the goal is to search exhaustively for all possible off-target sites for a given candidate, the “-very-sensitive” alignment parameter set emphasizing sensitivity is used. Next, the pairwise sequence alignments identified are reconstructed from the SAM^38^ alignment result using sam2pairwise^39^. Finally, a machine learning model is used to predict the likelihood of a duplex occurring between the probe candidate and each off-target site, and the predictions are summed into an off-target score for each candidate (**Fig. 2A**).

**Figure 1.**
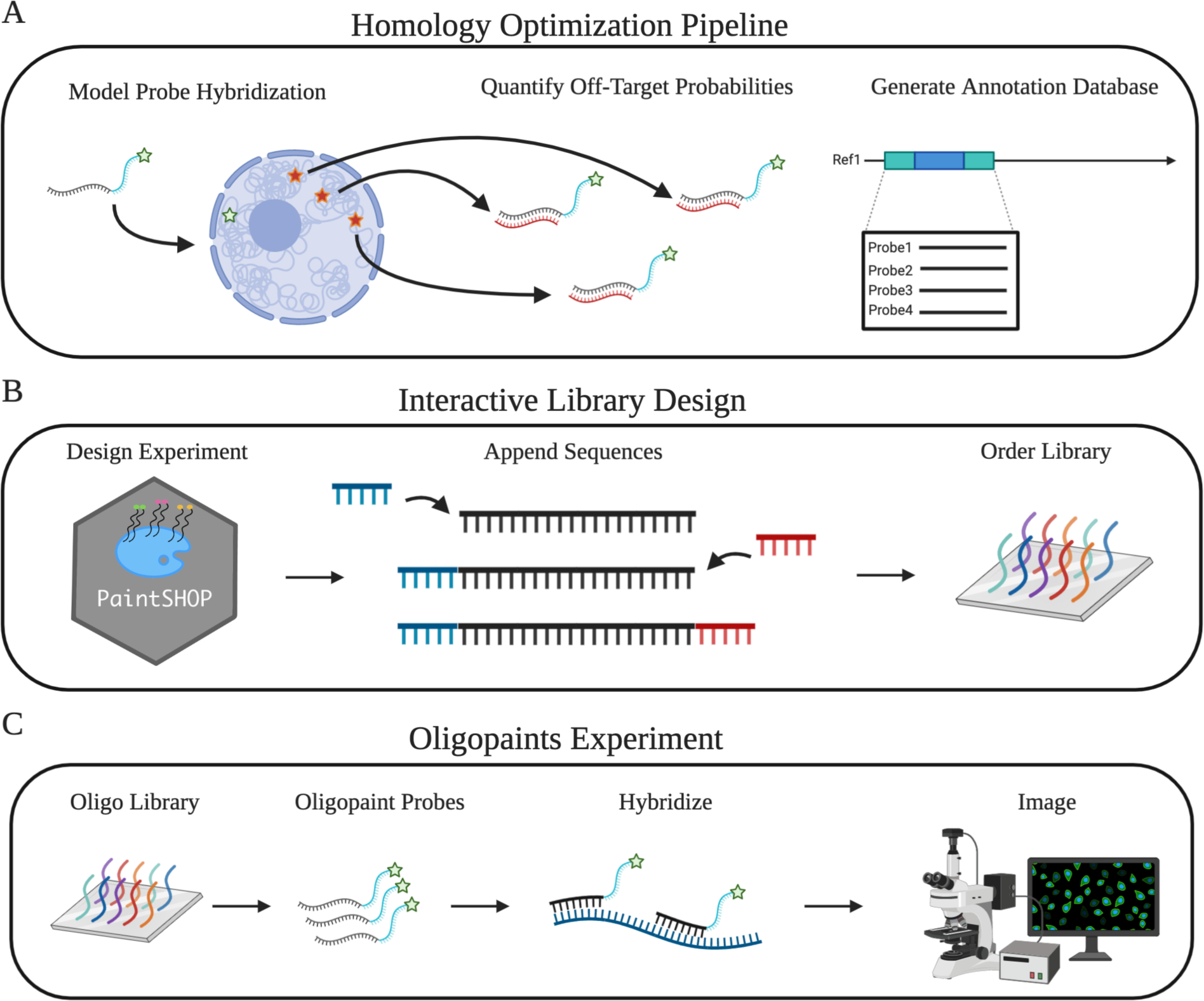
Overview of the PaintSHOP workflow. (A) Diagram of the homology optimization pipeline and creation of probe-transcriptome database. (B) Overview of the functionality of the web application provided to interact with PaintSHOP. (C) Schematic of the Oligopaints protocol.

**Figure 2.**
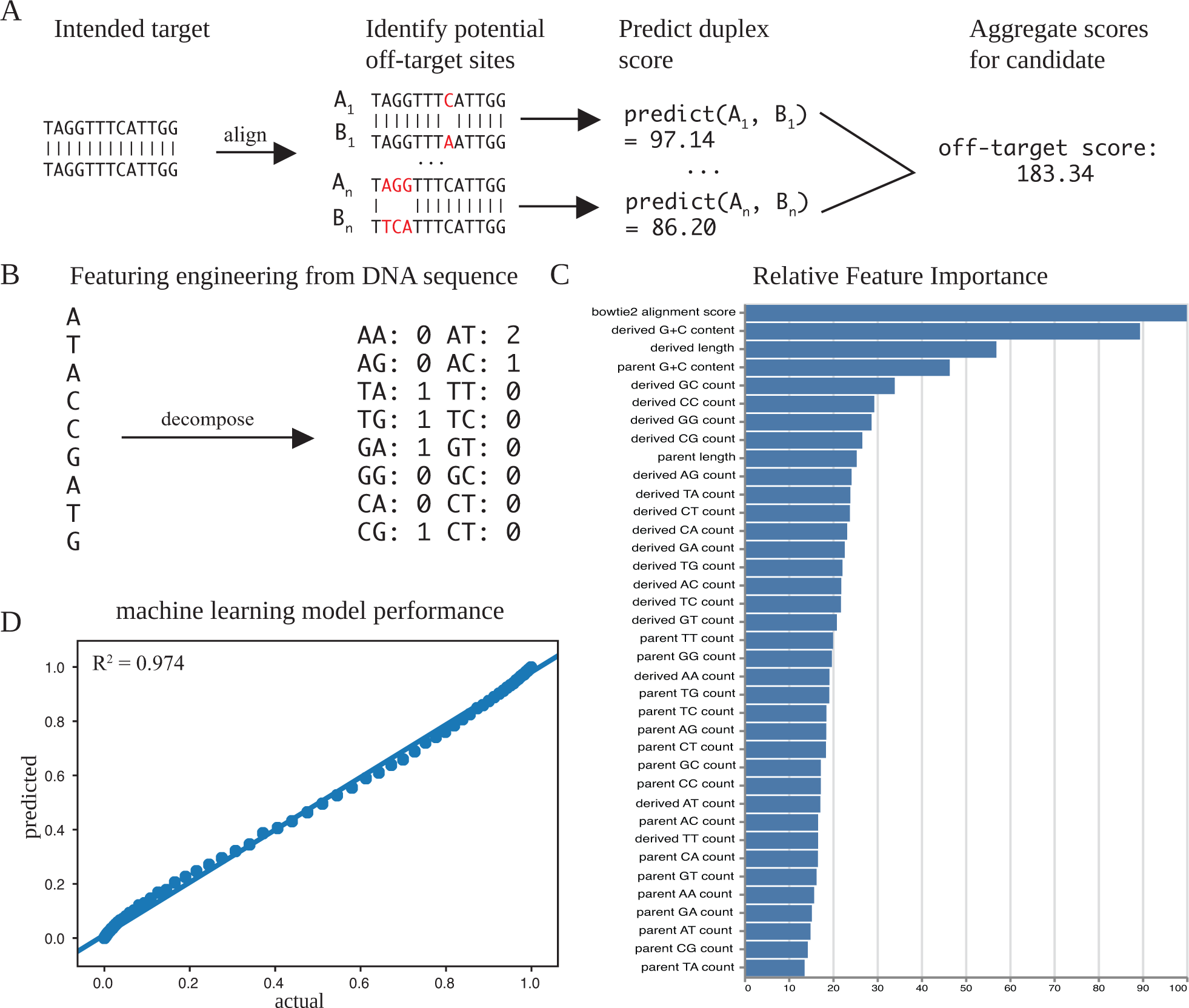
The homology optimization pipeline. (A) Diagram of the machine learning pipeline used to generate off-target scores for probe candidates. (B) Illustration of how dinucleotide counts are generated as numerical features of sequences as input for model prediction. (C) Ranked bar chart showing the relative feature importance of features incorporated in the predictive model. (D) A binned scatterplot (100 bins) of the model predictions versus the actual duplexing probabilities, with the least squares regression line shown.

The machine learning model works by approximating the duplexing probability generated by NUPACK pairwise test-tube simulations based on numerical features computed from the pairwise alignments (**Methods, Fig. 2B and C**). The underlying predictor is an XGBoost^40^ Regressor, which was selected after empirically evaluating a wide variety of possible models and hyperparameters using the TPOT^41,42^ genetic algorithm. The model predictions achieved a root-mean-square error (RMSE) score (**Methods**) of 0.0657, and the R^2^ score between actual and predicted values on the test set (n = 101,704) was 0.974 (**Fig. 2D**). Importantly, the ability to accurately predict the pairwise duplexing probability without directly computing NUPACK simulations is what makes the large-scale modeling of off-target binding computationally feasible. For example, as part of this work we computed >140,000,000 duplex predictions in less than one day, which would have taken more than two months with direct computation.

### Mining Novel Probe Sets

In order to perform an oligo-based FISH assay, it is necessary to identify the regions within your target sequence of interest that are most amenable to successful hybridization. These regions can be identified computationally by searching for as many candidates as possible that satisfy criteria necessary for successful FISH such as having relatively uniform length, GC-content, and *T*_*m*_, while screening for negative characteristics such as homopolymeric runs and repetitive sequences. Leveraging existing tools for this search process^33^, we performed a systematic search for probe sequence parameters that increased the total number of candidates identified while maintaining the quality and uniformity of the candidates. This search culminated in the creation of the “newBalance” probe sets for the hg38, hg19, mm10, mm9, dm6, and ce11 genome assemblies. The newBalance probe sets have a new minimum and maximum probe length window of 30-37, and can include repeat-masked^43^ bases .These changes allow users to optionally include these sequences in their design if it is necessary to increase the number of possible probe candidates covering their target of interest, which can be particularly valuable for RNA FISH where repetitive sequences are less of a concern and finding enough quality probes can be challenging.

To our knowledge, the newBalance probe sets constitute the largest sets of probe candidates generated to date. For hg38, the newBalance probe set contains over 41 million candidates, which is 2.7 times more probes than the largest previously published probe set for the assembly^33^ (**Fig. 3**). For hg19, newBalance has 1.38 times more probes than the second largest set^16^ (**Fig. 3**). In addition to a substantial increase in the number of candidates, the newBalance probe set provides a greater number of probes that pass through the homology optimization pipeline compared to previous probe sets (**Fig. 3**). We have augmented these *in silico* analyses by validating that newBalance probes behave as expected *in situ* by targeting the *ADAMTS5* mRNA in human kidney mesangial cells (**Supplementary Fig. S1**).

**Figure 3.**
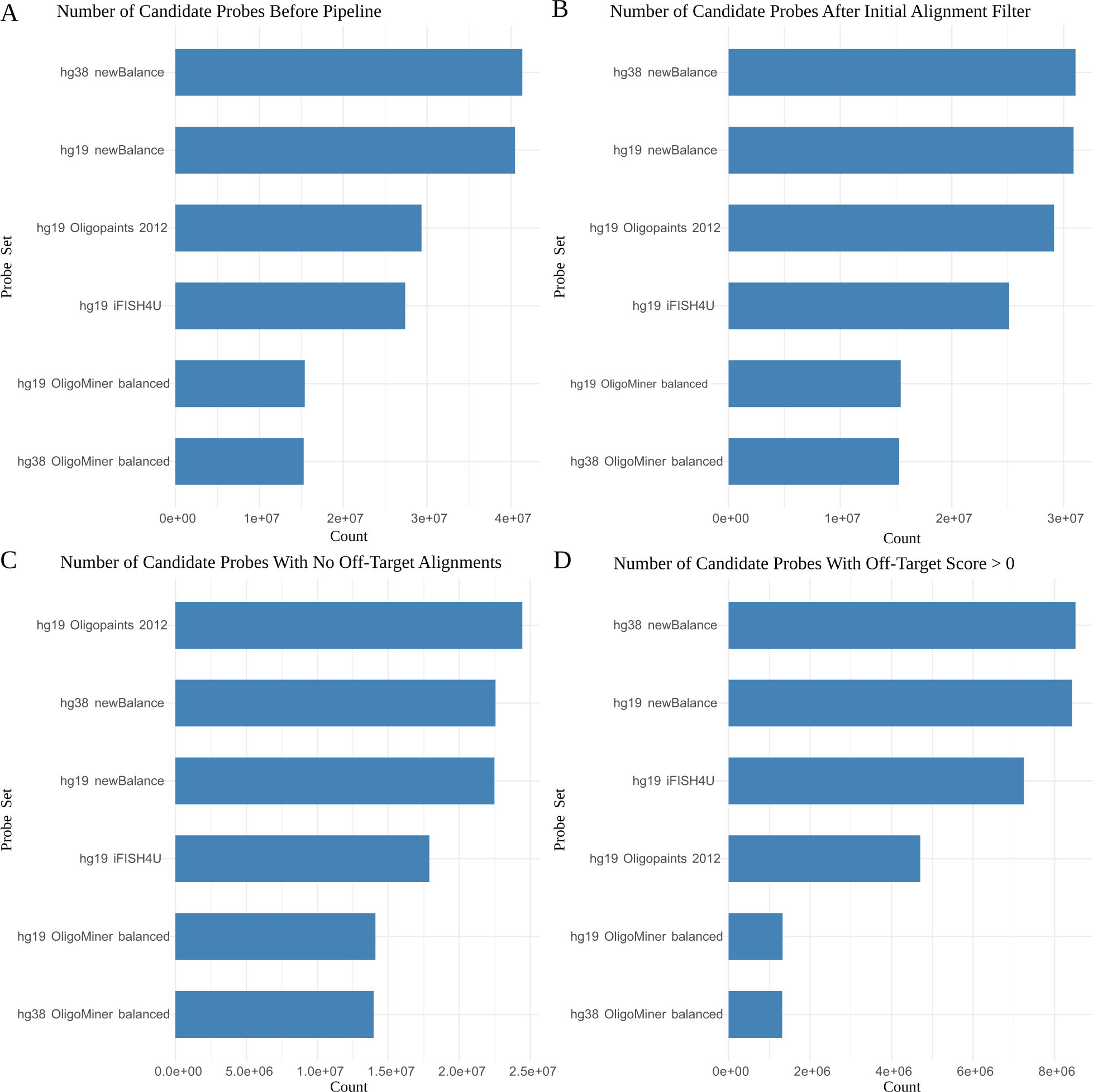
Counts of probes for each human probe set included in PaintSHOP. Counts are shown for the numbers of candidates before any downstream processing was performed (A), the number of candidates after filtering for probes with greater than 100 alignments (B), the numbers of remaining probes with no alignments (C) and the with between 1 and 100 off-target alignments (D).

### Transcriptome Intersection

Tools now exist to search across a genome for probes that can be used for oligo-based FISH^33,34^. While these tools efficiently generate genome-scale sets of oligo probes, to the best of our knowledge no comprehensive database exists to connect the coordinates of the discovered probes with the location of reference annotations such as RefSeq. We set out to create this resource in order to greatly reduce the computational difficulty of retrieving probes for single-molecule FISH^12^ (smFISH) experiments with multiple targets. Leveraging the ability to perform fast intersection operations on genomic coordinates with BEDTools^44^, we developed a flexible approach to intersect any probe set stored in Browser Extendible Data (BED) format^45^ with any annotation set for the assembly. This database enables the retrieval of the probes for an arbitrary number of targets with a simple lookup operation rather than performing a large number of manual intersections to retrieve probes for each target.

The ability to create probe-annotation databases enabled us to compare distributions of annotation coverage between probe sets (**Fig. 4A and Supplementary Fig. S2**). For the hg19 assembly, the average number of probes per transcript was 41.48 for the Oligopaints probe set from 2012^16^, 34.54 for the OligoMiner^33^ ‘balance’ probe set, 32.44 for the iFISH4U^34^ “full 40-mer” probe set, and 50.69 for the newBalance probe set introduced here. Additionally, for the newBalance probe set, nearly all (98.8%) of the hg19 annotations were targeted by at least one probe, and 85% of annotations were targeted by more than 15 probes (**Fig. 4B and Supplementary Fig. S2**), providing a considerable increase in probe coverage relative to previous probe sets. Importantly, our web application is agnostic to which probe set is used, thus allowing users to harness our newly developed newBalance probes, a large number existing of publicly available probe sets, and any additional probe sets that may be released at a future date.

**Figure 4.**
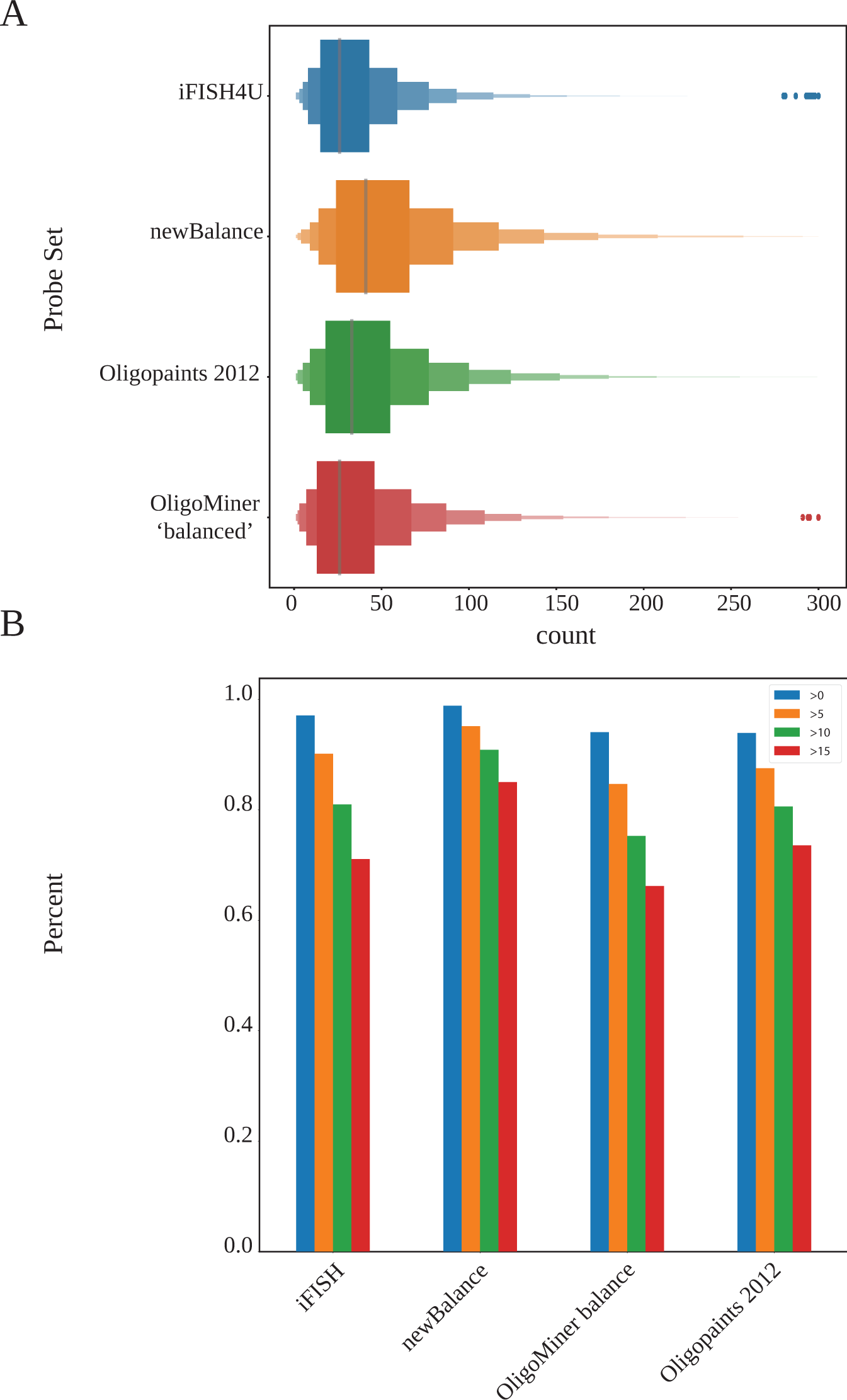
(A) Letter-value plots (boxplots with additional quantiles) of the distribution of the number of probes per transcript for all RefSeq annotations for the hg19 genome assembly. Probe sets shown are the iFISH4U “Full 40-mer” set (n = 2,219,982), the hg19 newBalance set (n = 3,531,893), the hg19 probe set from the original 2012 Oligopaints publication (n = 2,745,963), and the hg19 ‘balanced’ probe set generated by OligoMiner (n = 2,290,242). (B) The respective fraction of RefSeq annotations covered by greater than 0, 5, 10, and 15 probes after intersection with each genome-scale collection of hg19 oligo probes.

### Web-Based Interactive Design Application

In order to successfully perform an oligo-based RNA or DNA FISH experiment, a researcher must: 1) retrieve the probes covering their target(s); 2) ensure the probe sequences have the desired strand orientation; 3) consider trimming or balancing the number of probes per target in their set; 4) append the necessary primers and barcode sequences for their experimental design; 5) appropriately format an order file for a provider of the oligos. There has been no comprehensive resource or tool that was built to efficiently solve these design challenges for oligo-based FISH. As a result, computational probe design has remained a tedious problem for researchers that have enough command line experience to piece together approaches on their own that requires a substantial manual input and has further served as a barrier for many researchers to adopt the technology altogether. To address this bottleneck, we have built the web-based PaintSHOP interactive design application, which enables researchers to effectively and easily design probes for their oligo-based FISH experiments using an interactive, graphical web interface (**Supplementary Fig. S3**).

The PaintSHOP web application is designed to be modular and flexible, enabling a researcher to use only the features required for their experiment. Use cases can range from simply retrieving probes for a single RNA or DNA target to designing multi-target experiments with complex codebooks^22^. PaintSHOP is designed for users to be able to retrieve probes for their target, to optimize their set by adding or removing probes if necessary, to append primer and barcode sequences, and to generate an order file. For retrieving probes, users have the choice between two approaches: 1) RNA probe design and 2) DNA probe design. The RNA probe design option allows a user to either manually enter a list of RefSeq annotations or upload a file of annotations, and returns the probes that cover the inputted targets. The DNA design option accepts BED^45^ coordinates (chromosome, start coordinate, stop coordinate) either entered manually or uploaded from a file (**Supplementary Fig. S4**).

Once users have retrieved the probes for their target(s), they have the option to use several features to optimize the probe set returned. One crucial feature is the ability to tune several probe specificity parameters, enabling precise control over the inherent tradeoff between coverage and specificity (**Supplementary Fig. S5**). Previous approaches to modeling probe specificity have consisted of using binary classifiers to stringently filter candidates with any likelihood of off-target hybridization, resulting in highly specific probes at the cost of decreasing the number of probes, making the targeting of certain transcripts difficult or impossible^33^. Using the Homology Optimization Pipeline, we have generated a quantitative prediction of both on-target and off-target binding for every candidate probe in all probe sets in our application, making it possible to directly compare the predicted specificities of individual probes. Within the user interface, researchers can use these predictions to tune their probe sets, finding the balance between probe specificity and coverage that is the most optimal for their use case. Additionally, users can tune specificity based on the *k*-mer enrichment in the probes returned, and the inclusion or exclusion of repeat-masked^46^ sequences in their probes. Together, these parameters enable users to interactively explore how specificity scores impact the number of probes covering their targets through a dynamically updating interface.

While PaintSHOP enables control of specificity parameters to selectively increase target coverage, another common scenario in probe design is that there are too many probes to start with. An excess of probes can lead to an unbalanced set where some targets have far more probes than others and can result in unnecessarily expensive reagent costs. In order to remedy this, PaintSHOP has a “trim” feature, which allows a user to specify a desired number of probes per target and balances the set returned accordingly, with the probes predicted to be the most specific being kept. In certain cases, a researcher may have too few probes for some targets, and too many probes for others. For this scenario, PaintSHOP harnesses the quantitative specificity predictions to provide a “balance” feature, which selectively relaxes specificity parameters in order to cover targets with too few probes and trims targets with too many probes, resulting in a balanced probe set with a uniform number of probes for multiple targets.

A core advantage of oligo-based FISH is the precise control over the composition of the probe sequences that it provides, allowing for the incorporation of primer sequences and barcodes. For example, the Oligopaints^16^ technology requires the addition of PCR primer sequences to the 5’ and 3’ ends of the sequence homologous to the FISH target. The incorporation of primers enables the amplification of ssDNA oligo probes from the oligo library. Additionally, it is possible to incorporate region specific primers, allowing for more advanced imaging experiments such as “chromosome tracing”^47^ via sequential hybridization. In similar fashion to DNA FISH, advanced RNA FISH methods require the incorporation of multiple sequences in addition to the region homologous to the target into the final probe sequences. For example, spatial transcriptomics technologies such as MERFISH^22^ and seqFISH+^48^ require the addition of “barcode” sequences and “readout” sequences in order to perform more complicated experiments with many targets requiring successive hybridization. Similarly, SABER^49^, a molecular toolkit for FISH signal amplification and sequential hybridization, requires a Primer Exchange Reaction^50^ (PER) primer to be appended to the 3’ end of an oligo probe.

In order to accommodate a wide variety of oligo FISH technologies, PaintSHOP includes a flexible user interface for performing appending operations (**Fig. 5**). Through the interface it is possible to append up to three sequences to both the 5’ and 3’ end of each probe. For each sequence appended, the user can choose from a variety of encoding schemes. In the simplest case, a selected sequence can be appended to all probes in a given probe set (**Fig. 5A**). For example, a researcher can add the same 5’ primer to each probe in the set. PaintSHOP also allows a user to append a unique sequence to the probes for each target in a set (**Fig. 5B**). Using the same example, this would mean that a unique 5’ primer would be appended to the probes for each target in a set. To add addition flexibility, users can also add multiple sequences to a single position per target (**Fig. 5C**), or specify an entirely custom configuration. It is possible to quickly use PaintSHOP sequences provided for each position, or to upload a custom set of sequences to append. Collectively, the features in the flexible PaintSHOP appending interface can support a wide variety of oligo FISH technologies and experimental designs.

**Figure 5.**
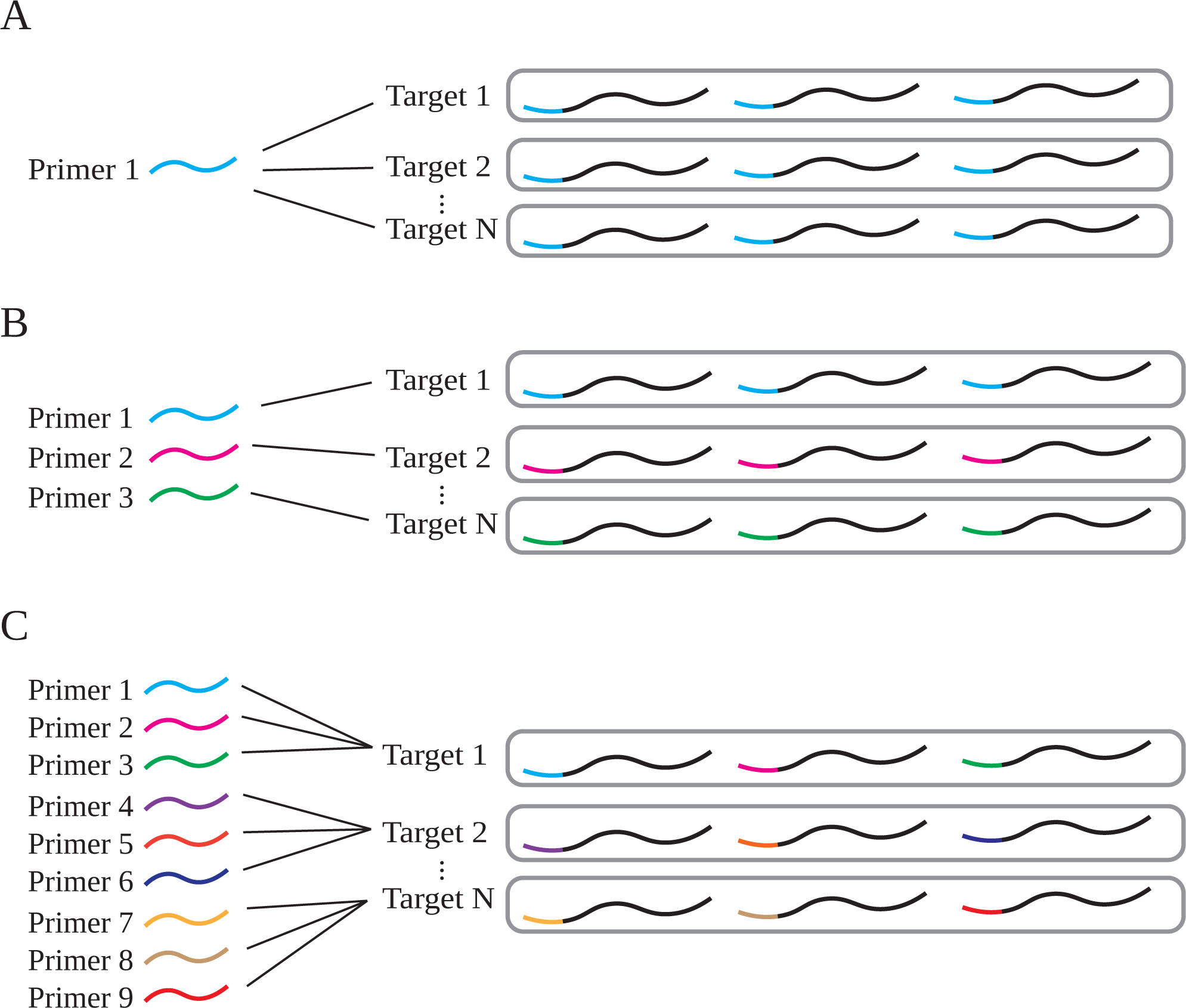
Schematic overview of possible appending schemes. (A) Appending the same primer to all probes for all targets in a set. (B) Appending a unique primer to all probes for each target in a set. (C) Appending a unique primer to each probe for each target in a set.

In addition to the general appending functionality, PaintSHOP provides built-in support for appending bridge sequences according to a MERFISH^22^ codebook. With this feature, users can upload a set of MHD4 16-bit barcodes to use with their RNA FISH targets. PaintSHOP automatically generates valid MERFISH probe sets by parsing the barcodes provided and handling the incorporation of the encoded bridge sequences into the probe sequences for each target. To demonstrate this feature, PaintSHOP was used with a set of 90 RNA FISH targets and barcodes (**Supplementary Files 1 and 2**) and a set of 16 readout sequences (**Supplementary File 3**) to create an order file for a MERFISH experiment (**Supplementary File 4**). Using the hg38 newBalance probe set with default PaintSHOP parameters, the targets had an average of 65.9 probes covering them (**Supplementary Fig. S6**). Additionally, as the targets for MERFISH and other highly multiplexed RNA FISH experiments are often chosen based on single-cell RNA sequencing datasets that do not generally have the ability to resolve the specific isoform(s) that map to a given cell, we have introduced ‘isoform flattened’ versions of the RefSeq annotations for the ce11, dm6, mm10, hg19, and hg38 assemblies. These ‘isoform flattened’ annotation sets prioritize shared exonic sequence between isoforms (**Methods**) in order to maximize the chance of detection and only modestly reduce the coverage of the transcriptome when used for probe intersects (**Supplementary Fig. S7**). Collectively, these new resources will streamline the design and practical implementation of spatial transcriptomic experiments.

The final core feature of the PaintSHOP web application is the download functionality provided. Once a researcher has taken advantage of the features necessary for the design of their FISH experiment, they can freely download all the information necessary for a successful order of their designed library. Additionally, we provide several optional download files that promote reproducibility and clear documentation of design decisions made and primers used. By providing features for probe retrieval, set balancing and trimming, sequence appending, and the free download of completed designs, PaintSHOP aims to be the first comprehensive resource for the design of complex oligo FISH experiments.

## Discussion

PaintSHOP is a freely available computational framework that enables the interactive design of transcriptome- and genome-scale oligo-based FISH experiments. PaintSHOP consists of a scalable machine learning pipeline for the thermodynamic modeling of probe specificity, and a dynamic web application for probe retrieval, library design, and the creation of complete order files. Leveraging the predictions made by the machine learning pipeline, the web application provides substantial control over parameters that impact the coverage of FISH targets, providing the flexibility needed for designing probe sets against targets that have fewer optimal probes to start. In addition to the introduction of a new pipeline and web resource, we have developed newBalance probe sets by optimizing our previously reported approach for genome scale probe mining^33^ to increase genome-wide coverage. The newBalance probe sets for the human, mouse, fly, and worm genomes are freely available through PaintSHOP along with many other sets created by various technologies^33,34^. Our goal for these technologies and resources is to democratize the ability to design the libraries needed for a wide variety of oligo FISH experiments^16,22,48,49^ against any target in the genome or transcriptome. We anticipate that PaintSHOP will enable researchers to perform novel FISH experiments interrogating genome organization and the spatial location of gene activity.

## Methods

### Probe sets and Genome Assemblies

OligoMiner hg19 and hg38 ‘balance’ probe sets were downloaded from yin.hms.harvard.edu/oligoMiner. The hg19 probe set from the original Oligopaints study^16^ was downloaded from oligopaints.hms.harvard.edu. The hg19 iFISH4U ‘full 40 mer’ probe set was downloaded from ifish4u.org. The ce11, dm6, hg19, and hg38 genome assemblies were downloaded with soft-masking from genome.ucsc.edu.

### Probe Mining Optimization

OligoMiner^33^ was downloaded from github.com/brianbeliveau/OligoMiner. The blockParse script was modified to search for probes in soft-masked genome sequences, and to report candidates in soft-masked regions with a special flag. The modified blockParse script was used to mine for probe candidates for each genome assembly with the parameters “-l 20 -L 60 -t 42 -T 47” to identify all possible probes between 20 and 60 nucleotides in length with a Tm between 42 and 47 degrees. A sliding window was used to identify the 8-nucleotide length window with the highest number of candidates. The candidates with the newly optimized settings and specially flagged candidates in soft-masked regions were termed the ‘newBalance’ probe sets. All probe mining was performed in a Python 2.7 Anaconda environment^51^ with the dependencies required for OligoMiner (Python 2.7, Biopython, scikit-learn) on the Department of Genome Science Central Cluster at the University of Washington.

### PaintSHOP Bridge Set Creation

A set of 1,500 G-depleted 46 nt DNA sequences were generated using Python with the following probabilities for incorporating each base: 0.33 for A, 0.33 for T, 0.33 for C, and 0.0 for G. The following substrings were excluded: “AAAA”, “TTTT”, and “CCC”. Each sequence had a maximum predicted *Tm* of 42°C33,52. Duplexing probabilities were computed for all pairwise combinations of bridge sequences and their reverse complements. 1,065 sequences with a >0.99 probability of on-target duplexing, a maximum off-target duplexing probability of <=0.015 and an average off-target duplexing probability of <=0.0006 were kept. Pairwise duplexing probabilities were computed for the remaining sequences to screen for potential dimerization between bridge sequences. All simulations were performed with the following FISH conditions: 42° C, 50% formamide, 0.390M sodium. 0.65°C per % (vol/vol) formamide was used to scale temperature values in thermodynamic calculations^33,52^. The 1,065 sequences were aligned to the hg38, mm10, dm6, and ce11 references genomes with Bowtie2^37^ using the “--very-sensitive-local” settings. The 818 sequences that aligned 0 times were screened for k-mer sequences against all four reference genomes using the OligoMiner^33^ kmerFilter.py utility with the settings “-m 18 -k 10”. The 800 remaining sequences were used as the new PaintSHOP bridge set.

### Machine Learning Model Development

Model construction was performing using the “probe-target” data set described originally in OligoMiner^33^. Briefly the data set consists of 406,814 pairs of “probe” and “target site” sequences. The “target sites” were generated using a combination of in-silico truncation, insertion, and point mutation of the “probe” sequences. The data set contains a Bowtie2^37^ alignment score and the thermodynamic duplexing probability computed using NUPACK 3.0^53–55^ for each sequence pair. The following numeric features were engineered to represent the key thermodynamic properties of the “probe” and “target site” sequences: length, GC-content, and dinucleotide counts. These features and the Bowtie2 alignment scores were used to build a machine learning model to predict the duplexing probability of the sequence pairs. The data set was randomly split into a training set and a testing set using scikit-learn^56^. Automatic model selection and hyperparameter optimization was performed using TPOT^41,42^. Negative mean squared error was used as the scoring function. After 10 generations with a population size of 100, TPOT converged on a XGBoost^40^ regressor. All model selection and hyperparameter optimization was performed using 5-fold cross-validation. The mean squared error (MSE)

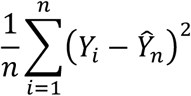

And root mean squared error

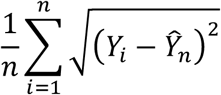

were computed for the test set, where *Ŷ*, is the predicted value, *Y* is the actual value, and *n* is the total number of samples. Least squares regression was also performed perform to evaluate the correlation between model predictions and actual values on the test set. Feature importance was calculated as the number of times a given feature was used to split the data across all trees.

### Pipeline Development

A pipeline to align mined “probe candidates” to the genome and compute off-target binding predictions was constructed using Snakemake^35^. The pipeline aligns “candidates” to their respective genome using Bowtie2^37^ with the settings “--very-sensitive-local -k 100”, returning up to 100 alignments. Pairwise alignments are reconstructed from the SAM format^38^ alignment results using sam2pairwise^39^. The XGBoost^40^ regressor is used to predict the duplexing probability of each pairwise alignment returned for all “probe candidates”. The “on-target score” is computed as

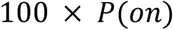

where *P*(*on*) is the duplexing probability at the intended “target” site, and the “off-target score” is computed as

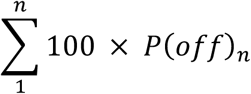

where *P*(*off*) is any alignment at a site other than the “target”, and *n* is the number of “off-target” alignments. Probabilities are scaled to the 0-100 range for user interpretability in the PaintSHOP web application. In addition to both scores, a *k*-mer statistic is computed. The number of times each 18-mer for a given candidate occurs within its respective genome is computed using Jellyfish^57^. The pipeline returns the occurrence count of the most frequently occurring 18-mer for each candidate.

### Creation of Collapsed Isoform Sets

GTF-format RefSeq annotation information for the ce11, dm6, mm10, hg19, and hg38 genome assemblies was obtained from the UCSC genome browser^45^. A custom Python script *iso-flatten.py* was created to identify genome intervals shared by all transcript isoforms of a given gene using information obtained from the ‘Coverage’ function of BEDTools^44^ called using pybedtools^58^. Genes for which no shared sequence exists among all the isoforms instead had the shared interval(s) shared by the greatest number of isoforms returned. The *iso-flatten*.*py* script is publicly available at: https://github.com/beliveau-lab/PaintSHOP_pipeline/tree/master/utilities.

### User Interface

A web application for interactive probe design was built using the Shiny^59^ web framework for the R programming language^60^. The back end of the application consists of two databases and a server. One database consists of the pre-computed set intersection of all probes for a given assembly with all UCSC RefSeq annotations in the assembly. The set intersection is computed using BEDTools^44^. The other database consists of all probes returned from the Homology Optimization Pipeline. The front end of the application enables interactive access to both databases. Users can either retrieve the probes targeting a set of RefSeq annotations, or can retrieve probes from the full database using any genomic coordinate in their assembly of interest. The front end also dynamically generates an interactive table for the user to view their probes, as well as a visualization of the distribution of probes per target using ggplot2^61^. An additional core feature implemented in the front end is the ability to append the sequences necessary for an oligo library, or a SABER^49^ experiment. All designs can be downloaded for use directly from the application.

### RNA SABER-FISH

The conditionally immortalized human mesangial cell line (K29Mes)^62^ was obtained from Dr. Moin Saleem (University of Bristol). Cells were cultured in RPMI-1640 medium supplemented with 10% FBS and ITS+ supplement. For propagation, the cells were grown at 33°C (permissive temperature). For experiments, cells were shifted to 37°C (non-permissive temperature) causing degradation of the temperature sensitive SV40 T-antigen and resulting in growth arrest. K29Mes cells were allowed to adhere to 22 x 22 #1.5 coverslips, then rinsed in 1x PBS, fixed in 4% (wt/vol) paraformaldehyde in 1x PBS for 10 minutes at room temperature, then rinsed in 1x PBS. Samples were then permeabilized in 1x PBS + 0.5% (vol/vol) Triton X-100 for 10 minutes at room temperature, then rinsed in 1x PBS + 0.1% (vol/vol) Tween-20. Samples were then transferred to 2x SSC + 1% (vol/vol) Tween-20 + 40% (vol/vol) formamide and incubated for 30 minutes at 43°C in a benchtop air incubator. Samples were then inverted onto parafilm square containing 80 µl of pre-warmed hybridization solution consisting of 2x SSC + 1% (vol/vol) Tween-20 + 40% (vol/vol) formamide + 10% (wt/vol) dextran sulfate and 80 µl of lyophilized product from a Primer Exchange Reaction (PER) reaction^49,50^ performed on a set of 105 oligo probes targeting the *ADAMTS5* mRNA (**Supplementary Table 2**). The *ADAMTS5* probe pool was purchased from Integrated DNA Technologies and was PER extended for 90 minutes at 37°C with a probe concentration of 1 µM and a hairpin h25.25^49^ concentration of 0.5 µM. Hybridization was allowed to proceed overnight (∼16 hours) at 43°C in a humidified chamber placed in a benchtop air incubator. Samples were then washed 2 times for 30 minutes each in 2x SSC + 1% (vol/vol) Tween-20 + 40% (vol/vol) formamide for 30 minutes at 43°C, and then twice for 5 minutes each in 2X SSC + 0.1% (vol/vol) Tween-20 at 43°C, and then twice for 5 minutes each in 1x PBS at room temperature. Samples were then inverted onto parafilm containing 100 µl of a secondary hybridization buffer consisting of 0.16x PBS + 8% (wt/vol) dextran sulfate + 0.04% (vol/vol) Tween-20 and an ATTO565-labeled p25*.25* secondary oligo^49^ at 0.4 µM and incubated for 30 minutes at 37°C in a benchtop air incubator. Samples were then washed twice for 5 minutes each in 1x PBS + 0.1% (vol/vol) Tween-20 at 37°C, then stained in 0.1 µg/ml DAPI in 1x PBS for 5 minutes at room temperature. Samples were then washed for 5 minutes in 1x PBS at room temperature, then inverted onto microscope slides containing ProLong Gold Antifade Mountant which cured overnight at room temperature prior to imaging. Images were captured using a Leica SP8X laser scanning confocal microscope using a 63x oil N.A. 1.40 Plan Apo objective lens controlled using Leica LASX Expert software. Images were processed in ImageJ + Fiji^63,64^ and Adobe Photoshop.

## Code and Data Availability

The source code for the PaintSHOP web application is available at https://github.com/beliveau-lab/PaintSHOP. The source code for the Homology Optimization Pipeline is available at https://github.com/beliveau-lab/PaintSHOP_pipeline. All genome-scale probe collections, primer sequences, bridge sequences, SABER-associated sequences, and transcriptome intersects hosted on paintshop.io are available to download from https://github.com/beliveau-lab/PaintSHOP_resources repository. All repositories are available under a MIT license

## Acknowledgements

The authors thank G. Nir, D. Shechner, A. Tsue, H. Nguyen, J.Y. Kishi, and J. Harke for helpful feedback during the beta testing phase of PaintSHOP, N. Peters for assistance with microscopy, the Genome Sciences IT team for technical assistance, and members of the Beliveau lab for feedback on the written manuscript. We also thank S. Lapan and E. West for productive discussions that inspired aspects of this work. This work was supported by a Damon Runyon Dale F. Frey Breakthrough Award (to B.J.B.) and the DiaCOMP consortium (19AU3987 to S.A. and B.J.B.). Imaging on the University of Washington W.M. Keck Center Lecia SP8X confocal microscopy was enabled by funding from the NIH (S10 OD016240). We would also like to thank the Paul G Allen Family Foundation for their support and encouragement through the Allen Institute for Brain Science.

