## Supplementary Information for "PaintSHOP enables the interactive design of transcriptome- and genome-scale oligonucleotide FISH experiments"

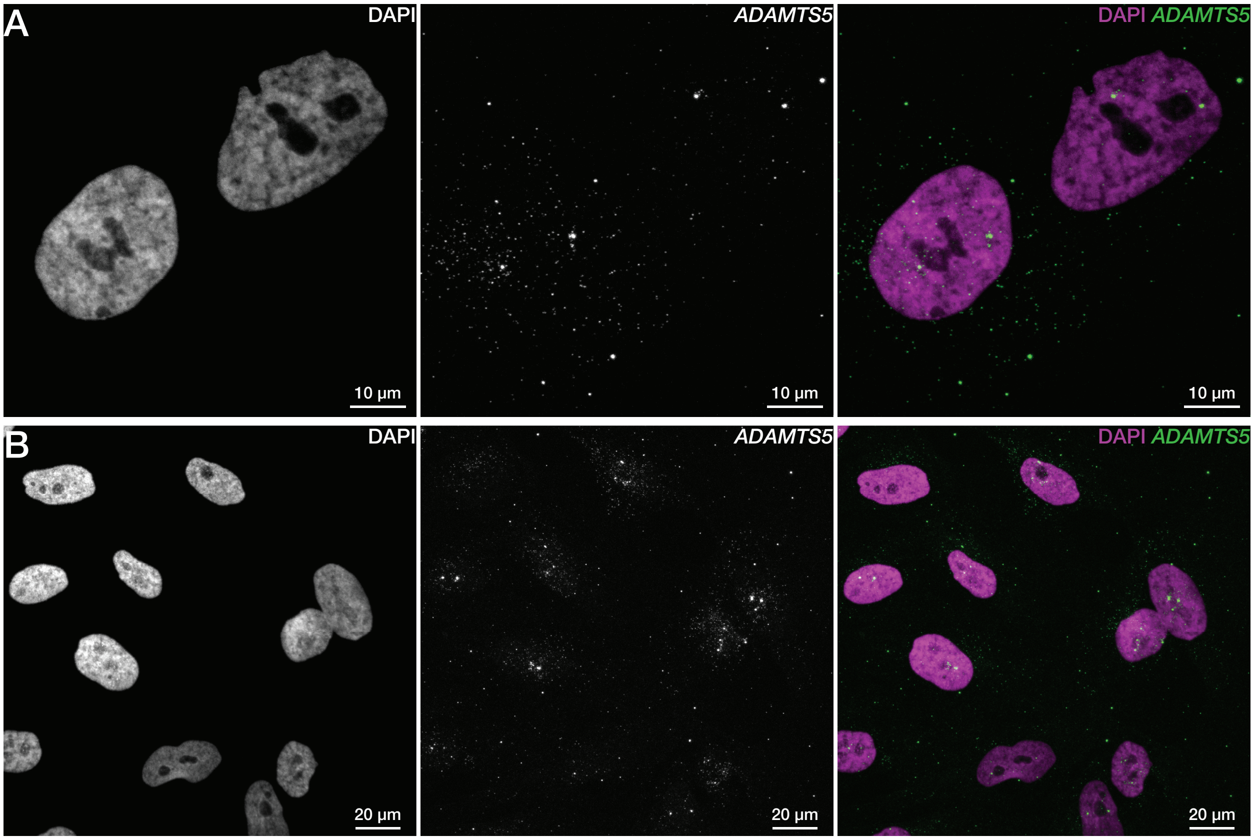

**Figure S1.** SABER-FISH was performed on the conditionally immortalized human mesangial cell line (K29Mes) using a set of 105 newBalance probe oligos targeting *ADAMTS5*. Images are maximum projections in Z. Also see Supplementary Table 2.

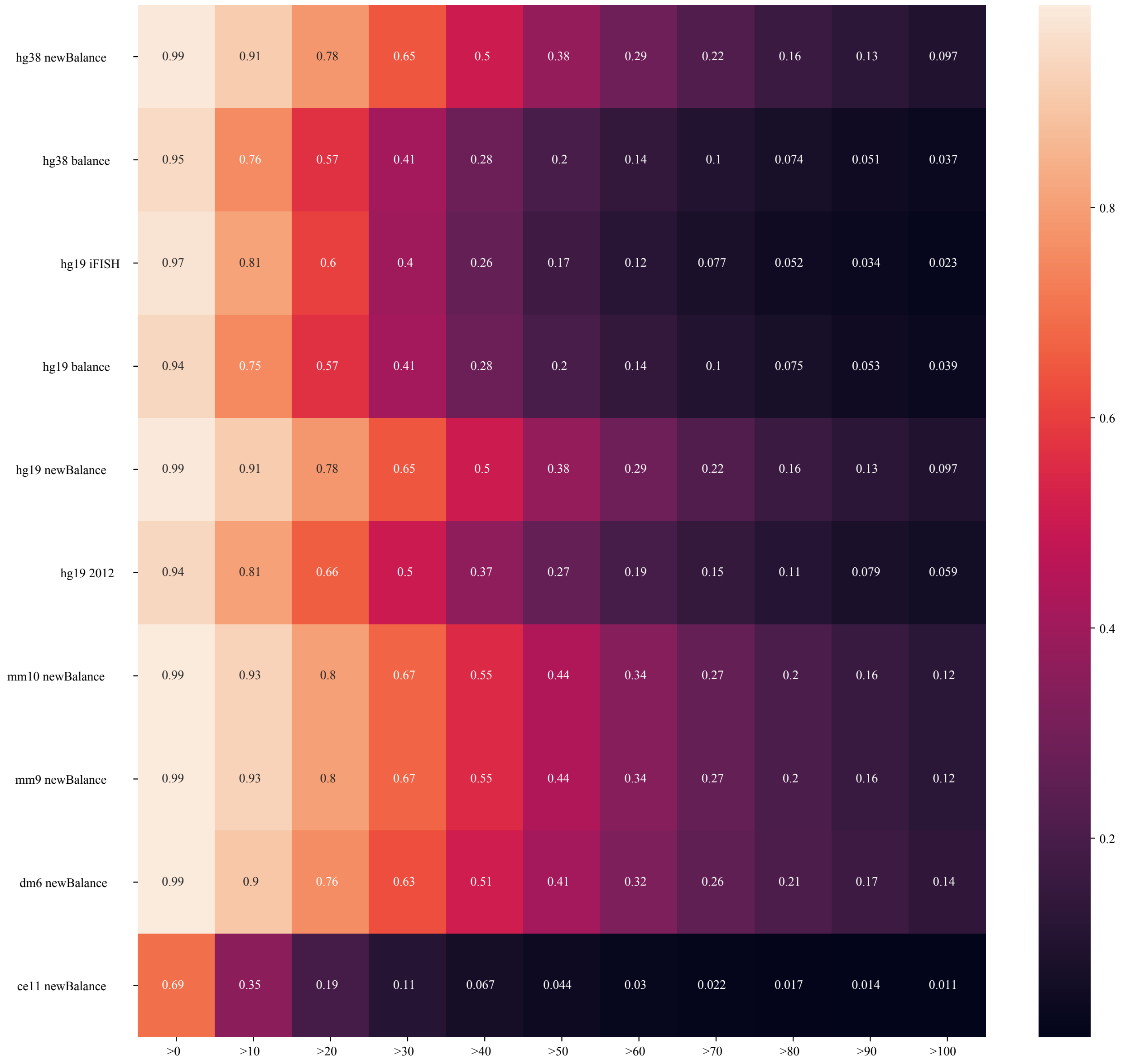

**Figure S2.** A heatmap of annotation probe coverage for all genome-scale probe sets included in PaintSHOP. Respective percentages of annotations with greater than 0, 10, 20, 30, 40, 50, 60, 70, 80, 90, and 100 probes that target them.

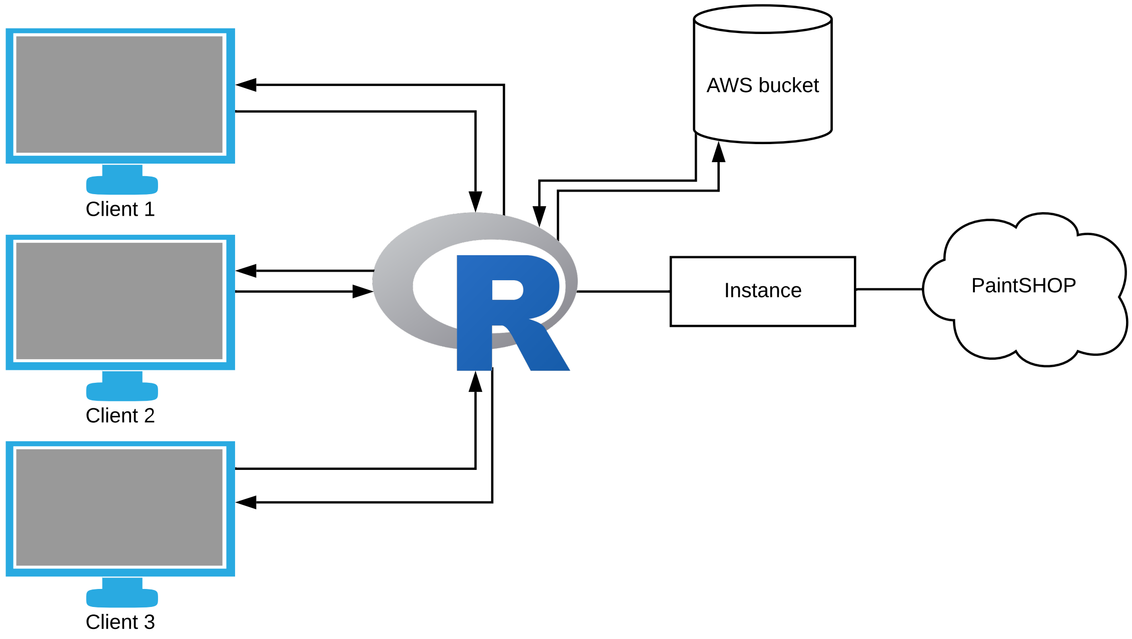

**Figure S3.** The architecture of the PaintSHOP application. The application follows a conventional client-server architecture. Using the shinyapps.io service, when a client connects to the web application, PaintSHOP initiates an application instance, which can run multiple R workers. An R worker can serve multiple clients. When a client submits a request for probes to the R worker, the worker retrieves the necessary data from an AWS bucket in the same region as the instance.

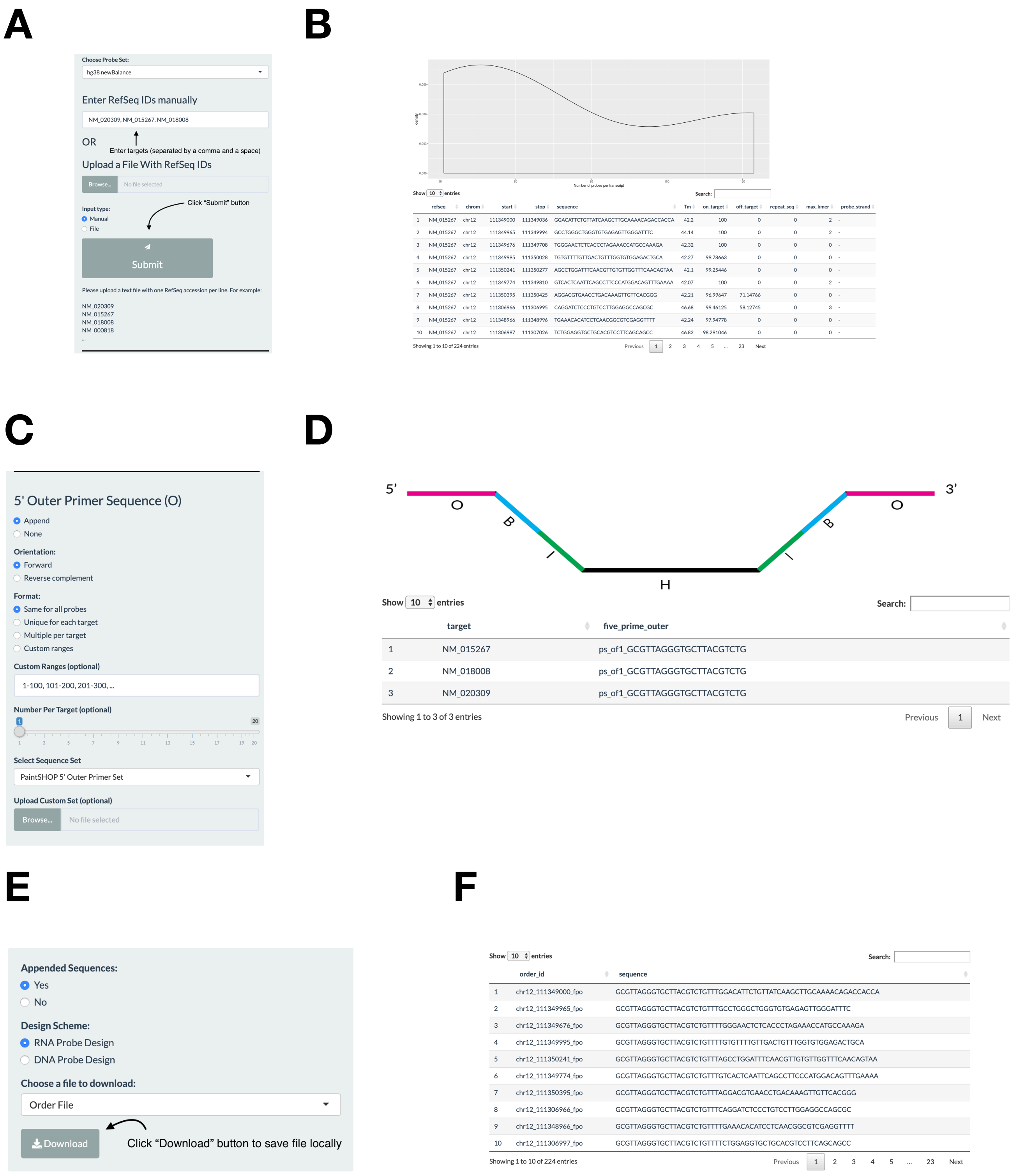
A
Choose Probe Set:
hg38 newBalance
Enter RefSeq IDs manually
NM_020309, NM_015267, NM_018008
OR
Enter targets (separated by a comma and a space)
Upload a File With RefSeq IDs
Browse...
No file selected
Click "Submit" button
Input type:
Manual
File
Submit
Please upload a text file with one RefSeq accession per line. For example:
NM_020309
NM_015267
NM_018008
NM_000818
...
B
Number of probes per transcript
Show 10 entries
Search:
refseq chrom start stop sequence Tm on_target off_target repeat_seq max_kmer probe_strand
1 NM_015267 chr12 111349000 111349036 GGACATTCTGTTATCAAGCTTGCAAAACAGACCACCA 42.2 100 0 0 2 -
2 NM_015267 chr12 111349965 111349994 GCCTGGGCTGGGTGTGAGAGTTGGGATTTC 44.14 100 0 0 2 -
3 NM_015267 chr12 111349676 111349708 TGGGAACTCTCACCCTAGAAACCATGCCAAAGA 42.32 100 0 0 0 -
4 NM_015267 chr12 111349995 111350028 TGTGTTTTGTTGACTGTTTGGTGTGGAGACTGCA 42.27 99.78663 0 0 0 -
5 NM_015267 chr12 111350241 111350277 AGCCTGGATTTCAACGTTGTGTTGGTTTCAACAGTAA 42.1 99.25446 0 0 0 -
6 NM_015267 chr12 111349774 111349810 GTCACTCAATTCAGCCTTCCCATGGACAGTTTGAAAA 42.07 100 0 0 2 -
7 NM_015267 chr12 111350395 111350425 AGGACGTGAACCTGACAAAGTTGTTCACGGG 42.21 96.99647 71.14766 0 0 -
8 NM_015267 chr12 111306966 111306995 CAGGATCTCCCTGTCCTTGGAGGCCAGCGC 46.68 99.46125 58.12745 0 3 -
9 NM_015267 chr12 111348966 111348996 TGAAACACATCCTCAACGGCGTCGAGGTTTT 42.24 97.94778 0 0 0 -
10 NM_015267 chr12 111306997 111307026 TCTGGAGGTGCTGCACGTCCTTCAGCAGCC 46.82 98.291046 0 0 0 -
Showing 1 to 10 of 224 entries
Previous 1 2 3 4 5 ... 23 Next
C
5' Outer Primer Sequence (O)
Append
None
Orientation:
Forward
Reverse complement
Format:
Same for all probes
Unique for each target
Multiple per target
Custom ranges
Custom Ranges (optional)
1-100, 101-200, 201-300, ...
Number Per Target (optional)
Select Sequence Set
PaintSHOP 5' Outer Primer Set
Upload Custom Set (optional)
Browse...
No file selected
D
5'
3'
O
B
H
B
O
Show 10 entries
Search:
target five_prime_outer
1 NM_015267 ps_of1_GCGTTAGGGTGCTTACGTCTG
2 NM_018008 ps_of1_GCGTTAGGGTGCTTACGTCTG
3 NM_020309 ps_of1_GCGTTAGGGTGCTTACGTCTG
Showing 1 to 3 of 3 entries
Previous 1 Next
E
Appended Sequences:
Yes
No
Design Scheme:
RNA Probe Design
DNA Probe Design
Choose a file to download:
Order File
Download
Click "Download" button to save file locally
F
Show 10 entries
Search:
order_id sequence
chr12_111349000_fpo GCGTTAGGGTGCTTACGTCTGTTTGGACATTCTGTTATCAAGCTTGCAAAACAGACCACCA chr12_111349965_fpo GCGTTAGGGTGCTTACGTCTGTTTGCCTGGGCTGGGTGTGAGAGTTGGGATTTC chr12_111349676_fpo GCGTTAGGGTGCTTACGTCTGTTTTGGGAACTCTCACCCTAGAAACCATGCCAAAGA chr12_111349995_fpo GCGTTAGGGTGCTTACGTCTGTTTTGTGTTTTGTTGACTGTTTGGTGTGGAGACTGCA chr12_111350241_fpo GCGTTAGGGTGCTTACGTCTGTTTAGCCTGGATTTCAACGTTGTGTTGGTTTCAACAGTAA chr12_111349774_fpo GCGTTAGGGTGCTTACGTCTGTTTGTCACTCAATTCAGCCTTCCCATGGACAGTTTGAAAA chr12_111350395_fpo GCGTTAGGGTGCTTACGTCTGTTTAGGACGTGAACCTGACAAAGTTGTTCACGGG chr12_111306966_fpo GCGTTAGGGTGCTTACGTCTGTTTCAGGATCTCCCTGTCCTTGGAGGCCAGCGC chr12_111348966_fpo GCGTTAGGGTGCTTACGTCTGTTTTGAAACACATCCTCAACGGCGTCGAGGTTTT chr12_111306997_fpo GCGTTAGGGTGCTTACGTCTGTTTTCTGGAGGTGCTGCACGTCCTTCAGCAGCC
Showing 1 to 10 of 224 entries
Previous 1 2 3 4 5 ... 23 Next

**Figure S4.** An example "RNA Probe Design" workflow. (A) The first step in the workflow consists of either manually entering targets separated by a comma and a space, or uploading a text file with a target on each line. Next, the user presses "Submit" to generate the probes. (B) An example of the graph and table returned describing the probe set that covers the targets entered. The table is searchable and resizable. (C) An example of parameter choices for appending a 5' outer (O) primer to the probes. Once a user presses the "Append" button, the sequences will be added to the probes. (D) The table returned once the sequences have been appended. The table shows which sequence has been added to each target in the set. The information in the given column is "[Primer set]_[primer type and number]_sequence". (E) An example of parameter choices for downloading probes from PaintSHOP. Once the user has selected whether they appended sequences and which type of design they did, the "Download" button can be used to save the files locally. (F) An example of the table showing the file selected to be downloaded. The order file consists of a column with a unique identifier for each sequence, and a column with the full probe sequences.

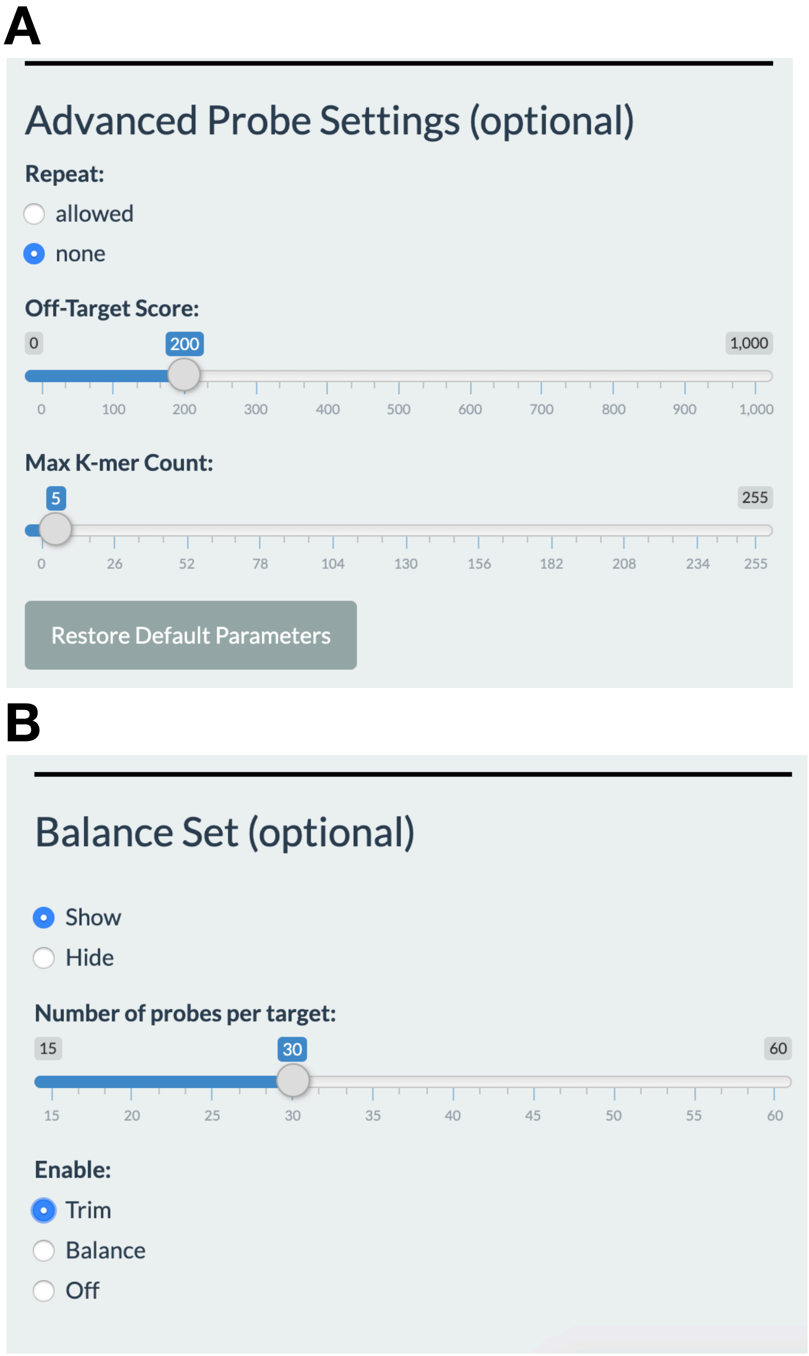

**Figure S5.** The optional advanced PaintSHOP settings and set balancing options. (A) The user can select whether repeat sequences are allowed in the probe sequences returned, and can tune the stringency both the off-target score and k-mer count used to filter probes with. (B) The set balancing options provided by PaintSHOP. The user can choose between the ‘trim’ option and ‘balance’ option described in the manuscript.

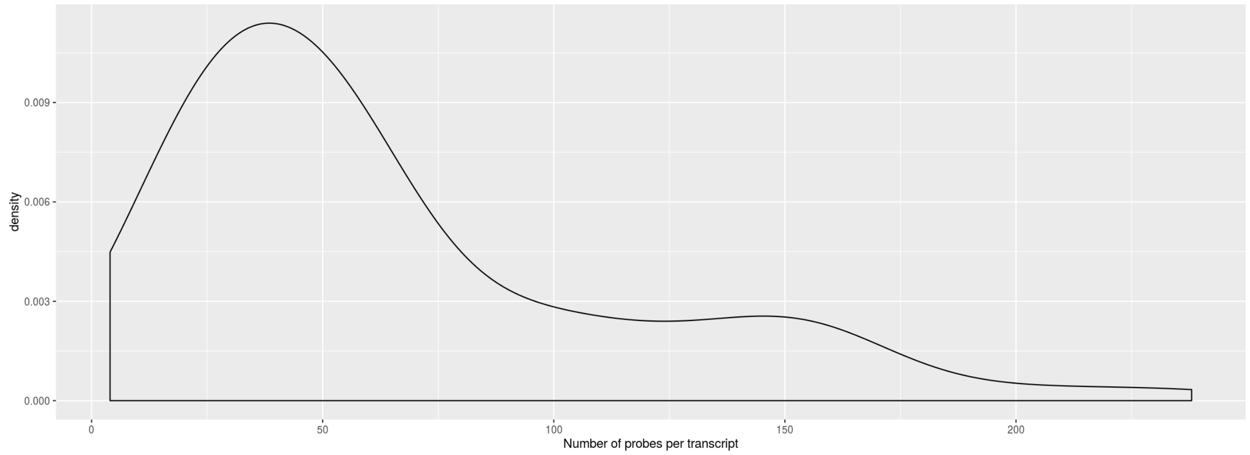

**Figure S6.** A density plot of the number of probes per transcript for the 90 RNA FISH targets in the MERFISH example set (Supplementary Files 1 and 2).

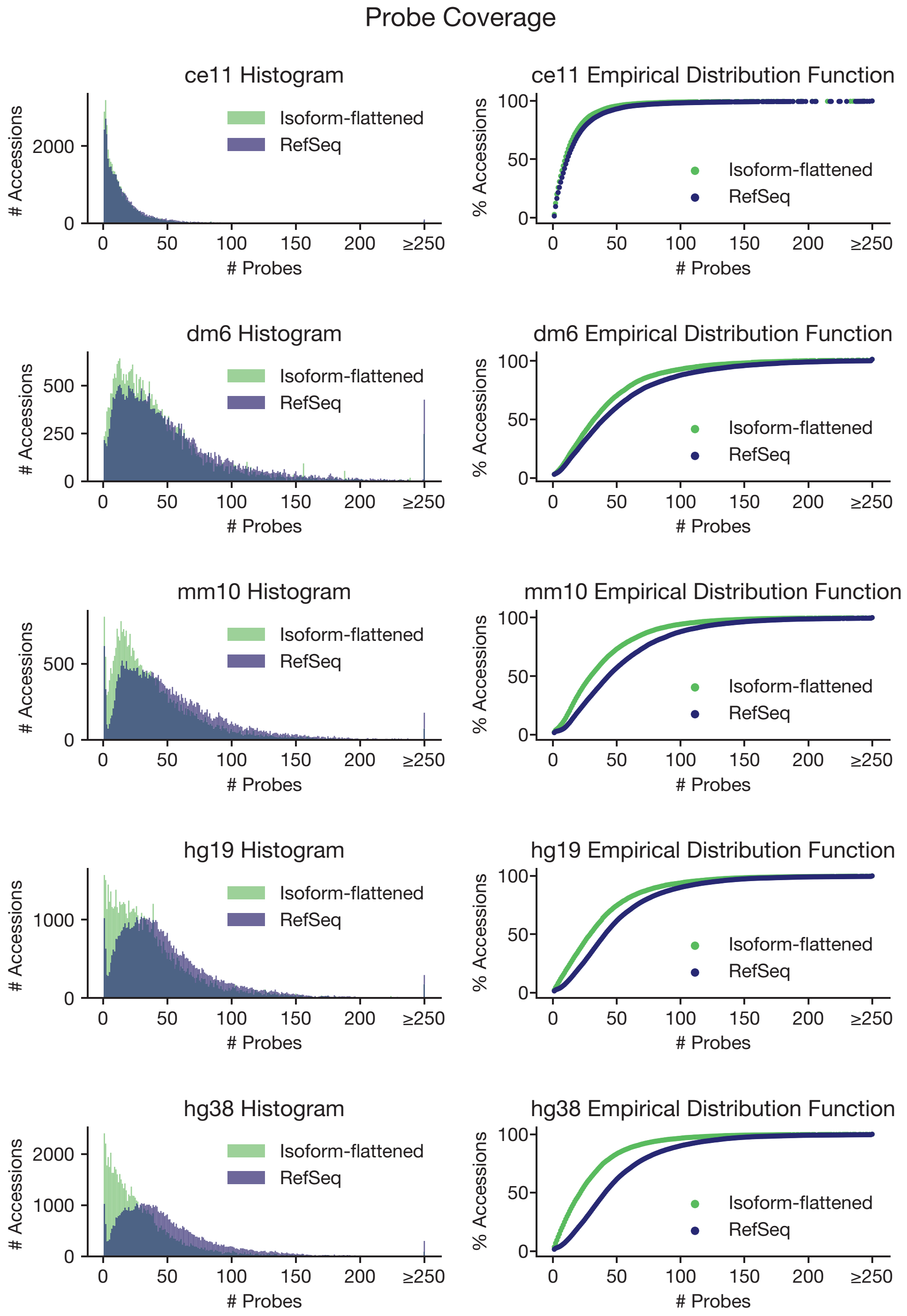

**Figure S7.** Histograms (left) and cumulative distribution plots (right) showing transcriptome-scale probe coverage when mapping to the standard UCSC RefSeq annotations (blue) and the "isoform-flattened" annotations introduced here (green). Only annotations present in both sets were considered for analysis.

| Set name | Number of sequences | Sequence length |
|---|---|---|
| Mateo et al. 2019 bridge sequences | 199 | 30 |
| PaintSHOP 3’ bridges | 532 | 46 |
| SABER 1x sequences | 50 | 10 |
| PaintSHOP outer forward primers | 10 | 21 |
| PaintSHOP bridge sequences | 1065 | 46 |
| SABER 2x sequences | 50 | 20 |
| PaintSHOP outer reverse primers | 10 | 21 |
| Xia et al. 2019 bridge sequences | 69 | 20 |
| PaintSHOP 5’ bridge sequences | 533 | 46 |
| MERFISH bridge sequences | 16 | 30 |
| PaintSHOP inner reverse primers | 74 | 21 |
| PaintSHOP inner forward primers | 74 | 21 |
| MERFISH primers | 318 | 20 |

**Table S1.** The number of sequences and sequence lengths for each set included in the PaintSHOP appending feature.

| | |
|---|---|
| chr21_26932943_saber | AGGGTTATTACAGTGACGATAGGCAAACTGCACTCCTTTACCAATAATAACCAATAATA |
| chr21_26932979_saber | TCCTCCACATGAGCGAGAACACTGGCCCCATTTACCAATAATAACCAATAATA |
| chr21_26933009_saber | GGATCCCCAAGATCCCCAGTTGCCATGGCTTTTACCAATAATAACCAATAATA |
| chr21_26934477_saber | TTTTCTTGGTTTTGTCCACACATTTGCCCTGCAGGTTTACCAATAATAACCAATAATA |
| chr21_26934512_saber | CAGATTCTCCCCTTTCCACAAGGCGTCCCTTTTACCAATAATAACCAATAATA |
| chr21_26934542_saber | TCCACCGCAGGCAGCTTCTTGGTCAGACAGTTTACCAATAATAACCAATAATA |
| chr21_26934572_saber | ACCATCTGGCCCTGGCGTACCACAGCACACTTTACCAATAATAACCAATAATA |
| chr21_26934602_saber | CACAGGCGAGCACAGACATCCATGCCGGGATTTACCAATAATAACCAATAATA |
| chr21_26934632_saber | CACACGGAGTACTCAGGCCCGAATGTCAGGTTTACCAATAATAACCAATAATA |
| chr21_26965555_saber | CATGGACGCGTCAGCCACCAGAAGCAGCTCTTTACCAATAATAACCAATAATA |
| chr21_26965654_saber | GGGCGAGAGAGCGGACTGGTCCAAGAGCTGTTTACCAATAATAACCAATAATA |
| chr21_26965787_saber | GCCTCGAAGCTGAAGCCCTCGCGGGTGTAGTTTACCAATAATAACCAATAATA |
| chr21_26965822_saber | CAGGATCCGTGCGGACCCATCCCCGTACACTTTACCAATAATAACCAATAATA |
| chr21_26965876_saber | GGGTCCGCGCAGCAGTGGCTTTAGGGTGTATTTACCAATAATAACCAATAATA |
| chr21_26965911_saber | CGTGCTTGACCGCGAAGAAGCCGTCGAGACTTTACCAATAATAACCAATAATA |
| chr21_26965941_saber | CCCCACAGAGGTCAAAGACAGCCAGAGAGCTTTACCAATAATAACCAATAATA |
| chr21_26965971_saber | GGGGACTACCGTCCACTGTGCCCCGATAGATTTACCAATAATAACCAATAATA |
| chr21_26966049_saber | GCACGAAGCCAGCAATGCCCACCGAACCATTTTACCAATAATAACCAATAATA |
| chr21_26966079_saber | CTCGCTCCAGGTCCAAGAGGAACCTCCGGCTTTACCAATAATAACCAATAATA |
| chr21_26966134_saber | CTTGCCGCCGCCGGAGTAGAGTTGGTCGATTTTACCAATAATAACCAATAATA |
| chr21_26966280_saber | CAGCAGTCGGAGGCTGCCCGGCTTTATCCTTTTACCAATAATAACCAATAATA |
| chr21_26966408_saber | GGAGGGGCTGCGCGACTGGGACTTTATGGGTATTTTTACCAATAATAACCAATAATA |

| | |
|---|---|
| chr21_26966556 _saber | GGCTGCGGTGCCAGCGTGCGAACTTTTCTTTTTACCAATAATAACCAATAATA |
| chr21_26966586 _saber | TGTTGCTTGGGAAAATGTTTGGATTCGTGCTCCATTTACCAATAATAACCAATAATA |
| chr21_26966668 _saber | AGTCGGATAGTGGAGATTCAGCAAATACGGGAAAAGGTTTACCAATAATAACCAATAATA |
| chr21_26966722 _saber | AAAACCACCAAATGCAGGCACGATCGCTGTTTTACCAATAATAACCAATAATA |
| chr21_26966752 _saber | TTCAAGAAAAGTAAGGAAAGGTACCGCGCACCGTTTACCAATAATAACCAATAATA |
| chr21_26966826 _saber | GCTGGACAAGCCGGCTGTGGAGGTTCCAAGTTTACCAATAATAACCAATAATA |
| chr21_26966856 _saber | CTCCGGGGCCCACTTCCTTTCTTATTTTGTGGTTTACCAATAATAACCAATAATA |
| chr21_26966888 _saber | GTTTCCTTGGCTCTGCTGGGACCAGGAGGATTTACCAATAATAACCAATAATA |
| chr21_26966984 _saber | GAGCTGGTGAAGCGGCGTGTCTGTGTCCCTTTTACCAATAATAACCAATAATA |
| chr21_26934662 _saber | TTGCACTGCTGGGTGGCATCGTAGGTCTGTTTTACCAATAATAACCAATAATA |
| chr21_26934692 _saber | CCTGGGAGTTCTTCGGGGCCCAGGATCTGCTTTACCAATAATAACCAATAATA |
| chr21_26943410 _saber | TGGCTGAAGTGCATTTGGACCAGGGCTTAGTTTACCAATAATAACCAATAATA |
| chr21_26943441 _saber | TGCATCAATGCTGGTAAGGATGGAAGACATTAAGCGCTTTACCAATAATAACCAATAATA |
| chr21_26943497 _saber | AGGTCTCTTCACAGAATTTGGAATCGTCATGGGAGAGTTTACCAATAATAACCAATAATA |
| chr21_26954742 _saber | TTTCGTGAGCCACAGTGAAGGCTGCGTGGATTTACCAATAATAACCAATAATA |
| chr21_26954772 _saber | GGCCATCGTCTTCAATCACAGCACAGCTGCTTTACCAATAATAACCAATAATA |
| chr21_26954802 _saber | GCTCTGGAGAACATATGGTCCCAACGTCTGCTTTACCAATAATAACCAATAATA |
| chr21_26954833 _saber | CATTCCCAGGGTGTCACATGAATGATGCCCATTTACCAATAATAACCAATAATA |
| chr21_26965298 _saber | AACAGGATAGCTGCATCGTAGTGCTCCTCATGGTTTACCAATAATAACCAATAATA |
| chr21_26965331 _saber | TCATCTCCCAGCTGGTTGTGTTGGTGCTGCTTTACCAATAATAACCAATAATA |
| chr21_26965361 _saber | CACTTGCAAAAGTTCTTGAGTGTGGTGGCAGCTTTACCAATAATAACCAATAATA |
| chr21_26965393 _saber | GTTCTTGCTCACTTCCAGGCTCTTGTCCTTGTCTTTACCAATAATAACCAATAATA |

| | |
|---|---|
| chr21_26965457_saber | CGGATGTGGTTCTCGATGCTAGCATGGCTGTTTACCAATAATAACCAATAATA |
| chr21_26932943_saber | AGGGTTATTACAGTGACGATAGGCAAACTGCACTCCTTTACCAATAATAACCAATAATA |
| chr21_26932979_saber | TCCTCCACATGAGCGAGAACACTGGCCCCATTTACCAATAATAACCAATAATA |
| chr21_26933009_saber | GGATCCCCAAGATCCCCAGTTGCCATGGCTTTTACCAATAATAACCAATAATA |
| chr21_26934477_saber | TTTTCTTGGTTTTGTCCACACATTTGCCCTGCAGGTTTACCAATAATAACCAATAATA |
| chr21_26934512_saber | CAGATTCTCCCCTTTCCACAAGGCGTCCCTTTTACCAATAATAACCAATAATA |
| chr21_26934542_saber | TCCACCGCAGGCAGCTTCTTGGTCAGACAGTTTACCAATAATAACCAATAATA |
| chr21_26934572_saber | ACCATCTGGCCCTGGCGTACCACAGCACACTTTACCAATAATAACCAATAATA |
| chr21_26934602_saber | CACAGGCGAGCACAGACATCCATGCCGGGATTTACCAATAATAACCAATAATA |
| chr21_26934632_saber | CACACGGAGTACTCAGGCCCGAATGTCAGGTTTACCAATAATAACCAATAATA |
| chr21_26965555_saber | CATGGACGCGTCAGCCACCAGAAGCAGCTCTTTACCAATAATAACCAATAATA |
| chr21_26965654_saber | GGGCGAGAGAGCGGACTGGTCCAAGAGCTGTTTACCAATAATAACCAATAATA |
| chr21_26965787_saber | GCCTCGAAGCTGAAGCCCTCGCGGGTGTAGTTTACCAATAATAACCAATAATA |
| chr21_26965822_saber | CAGGATCCGTGCGGACCCATCCCCGTACACTTTACCAATAATAACCAATAATA |
| chr21_26965876_saber | GGGTCCGCGCAGCAGTGGCTTTAGGGTGTATTTACCAATAATAACCAATAATA |
| chr21_26965911_saber | CGTGCTTGACCGCGAAGAAGCCGTCGAGACTTTACCAATAATAACCAATAATA |
| chr21_26965941_saber | CCCCACAGAGGTCAAAGACAGCCAGAGAGCTTTACCAATAATAACCAATAATA |
| chr21_26965971_saber | GGGGACTACCGTCCACTGTGCCCCGATAGATTTACCAATAATAACCAATAATA |
| chr21_26966049_saber | GCACGAAGCCAGCAATGCCCACCGAACCATTTTACCAATAATAACCAATAATA |
| chr21_26966079_saber | CTCGCTCCAGGTCCAAGAGGAACCTCCGGCTTTACCAATAATAACCAATAATA |
| chr21_26966134_saber | CTTGCCGCCGCCGGAGTAGAGTTGGTCGATTTTACCAATAATAACCAATAATA |
| chr21_26966280_saber | CAGCAGTCGGAGGCTGCCCGGCTTTATCCTTTTACCAATAATAACCAATAATA |

| | |
|---|---|
| chr21_26966408_saber | GGAGGGGCTGCGCGACTGGGACTTTATGGGTATTTTTTACCAATAATAACCAATAATA |
| chr21_26966556_saber | GGCTGCGGTGCCAGCGTGCGAACTTTTCTTTTTACCAATAATAACCAATAATA |
| chr21_26966586_saber | TGTTGCTTGGGAAAATGTTTGGATTCGTGCTCCATTTACCAATAATAACCAATAATA |
| chr21_26966668_saber | AGTCGGATAGTGGAGATTCAGCAAATACGGGAAAAGGTTTACCAATAATAACCAATAATA |
| chr21_26966722_saber | AAAACCACCAAATGCAGGCACGATCGCTGTTTTACCAATAATAACCAATAATA |
| chr21_26966752_saber | TTCAAGAAAAGTAAGGAAAGGTACCGCGCACCGTTTACCAATAATAACCAATAATA |
| chr21_26966826_saber | GCTGGACAAGCCGGCTGTGGAGGTTCCAAGTTTACCAATAATAACCAATAATA |
| chr21_26966856_saber | CTCCGGGGCCCACTTCCTTTCTTATTTTGTGGTTTACCAATAATAACCAATAATA |
| chr21_26932943_saber | AGGGTTATTACAGTGACGATAGGCAAACTGCACTCCTTTACCAATAATAACCAATAATA |
| chr21_26932979_saber | TCCTCCACATGAGCGAGAACACTGGCCCCATTTACCAATAATAACCAATAATA |
| chr21_26933009_saber | GGATCCCCAAGATCCCCAGTTGCCATGGCTTTTACCAATAATAACCAATAATA |
| chr21_26934477_saber | TTTTCTTGGTTTTGTCCACACATTTGCCCTGCAGGTTTACCAATAATAACCAATAATA |
| chr21_26934512_saber | CAGATTCTCCCCTTTCCACAAGGCGTCCCTTTTACCAATAATAACCAATAATA |
| chr21_26934542_saber | TCCACCGCAGGCAGCTTCTTGGTCAGACAGTTTACCAATAATAACCAATAATA |
| chr21_26934572_saber | ACCATCTGGCCCTGGCGTACCACAGCACACTTTACCAATAATAACCAATAATA |
| chr21_26934602_saber | CACAGGCGAGCACAGACATCCATGCCGGGATTTACCAATAATAACCAATAATA |
| chr21_26934632_saber | CACACGGAGTACTCAGGCCCGAATGTCAGGTTTACCAATAATAACCAATAATA |
| chr21_26965555_saber | CATGGACGCGTCAGCCACCAGAAGCAGCTCTTTACCAATAATAACCAATAATA |
| chr21_26965654_saber | GGGCGAGAGAGCGGACTGGTCCAAGAGCTGTTTACCAATAATAACCAATAATA |
| chr21_26965787_saber | GCCTCGAAGCTGAAGCCCTCGCGGGTGTAGTTTACCAATAATAACCAATAATA |
| chr21_26965822_saber | CAGGATCCGTGCGGACCCATCCCCGTACACTTTACCAATAATAACCAATAATA |
| chr21_26965876_saber | GGGTCCGCGCAGCAGTGGCTTTAGGGTGTATTTACCAATAATAACCAATAATA |

| | |
|---|---|
| chr21_26965911 _saber | CGTGCTTGACCGCGAAGAAGCCGTCGAGACTTTACCAATAATA ACCAATAATA |
| chr21_26965941 _saber | CCCCACAGAGGTCAAAGACAGCCAGAGAGCTTTACCAATAATA ACCAATAATA |
| chr21_26965971 _saber | GGGGACTACCGTCCACTGTGCCCCGATAGATTTACCAATAATA ACCAATAATA |
| chr21_26966049 _saber | GCACGAAGCCAGCAATGCCCACCGAACCATTTTACCAATAATA ACCAATAATA |
| chr21_26966079 _saber | CTCGCTCCAGGTCCAAGAGGAACCTCCGGCTTTACCAATAATA ACCAATAATA |
| chr21_26966134 _saber | CTTGCCGCCGCCGGAGTAGAGTTGGTCGATTTTACCAATAATA ACCAATAATA |
| chr21_26966280 _saber | CAGCAGTCGGAGGCTGCCCGGCTTTATCCTTTTACCAATAATA ACCAATAATA |
| chr21_26966408 _saber | GGAGGGGCTGCGCGACTGGGACTTTATGGGTATTTTTACCAA TAATAACCAATAATA |
| chr21_26966556 _saber | GGCTGCGGTGCCAGCGTGCGAACTTTTCTTTTTACCAATAATAA CCAATAATA |
| chr21_26966586 _saber | TGTTGCTTGGGAAAATGTTTGGATTCGTGCTCCATTTACCAATA ATAACCAATAATA |
| chr21_26966668 _saber | AGTCGGATAGTGGAGATTCAGCAAATACGGGAAAAGGTTTACC AATAATAACCAATAATA |
| chr21_26966722 _saber | AAAACCACCAAATGCAGGCACGATCGCTGTTTTACCAATAATAA CCAATAATA |
| chr21_26966752 _saber | TTCAAGAAAAGTAAGGAAAGGTACCGCGCACCGTTTACCAATA ATAACCAATAATA |
| chr21_26966826 _saber | GCTGGACAAGCCGGCTGTGGAGGTTCCAAGTTTACCAATAATA ACCAATAATA |
| chr21_26932943 _saber | AGGGTTATTACAGTGACGATAGGCAAACTGCACTCCTTTACCA ATAATAACCAATAATA |
| chr21_26932979 _saber | TCCTCCACATGAGCGAGAACACTGGCCCCATTTACCAATAATA ACCAATAATA |
| chr21_26933009 _saber | GGATCCCCAAGATCCCCAGTTGCCATGGCTTTTACCAATAATA ACCAATAATA |

**Table S2.** The RNA FISH probe set targeting the human *ADAMTS5* mRNA.
